# Optimizing motor decision-making through competition with opponents

**DOI:** 10.1101/405878

**Authors:** Keiji Ota, Mamoru Tanae, Kotaro Ishii, Ken Takiyama

**Affiliations:** Institute of Engineering, Tokyo University of Agriculture and Technology, Koganei, Tokyo, 184-8588, Japan; Department of Psychology, New York University, New York, NY, 10003, United States; Center for Neural Science, New York University, New York, NY, 10003, United States

## Abstract

Although optimal decision-making is essential for sports performance and fine motor control, it has been repeatedly confirmed that humans show a strong risk-seeking bias, selecting a risky strategy over an optimal solution. Despite such evidence, the ideal method to promote optimal decision-making remains unclear. Here, we propose that interactions with other people can influence motor decision-making and improve risk-seeking bias. We developed a competitive reaching game (a variant of the “chicken game”) in which aiming for greater rewards increased the risk of no reward and subjects competed for the total reward with their opponent. The game resembles situations in sports, such as a penalty kick in soccer, service in tennis, the strike zone in baseball, or take-off in ski jumping. In five different experiments, we demonstrated that, at the beginning of the competitive game, the subjects robustly switched their risk-seeking strategy to a risk-averse strategy. Following the reversal of the strategy, the subjects achieved optimal decision-making when competing with risk-averse opponents. This optimality was achieved by a non-linear influence of an opponent’s decisions on a subject’s decisions. These results suggest that interactions with others can alter human motor decision strategies and that competition with a risk-averse opponent is key for optimizing motor decision-making.

## Introduction

Optimal decision-making is indispensable for ideal performance in sports and fine motor control in everyday life. For example, selecting an appropriate trajectory for reaching a glass of water can lead to a low risk of spilling water, and likewise, finding a running path to easily pass through in rugby and deciding the best shot location in a tennis match can increase the chance of winning in a competition. Despite the importance of optimal decision-making, for over a decade, sub-optimal and overly risk-seeking behaviors have been reported in various motor decision tasks [1–11] (however, see also 12–14,15). Determining how to improve sub-optimal and risk-seeking decision-making behavior is crucial to enhance well-being in daily life and performance in sports. However, effective strategies to optimize human motor decision-making remain unknown.

One possible solution is to interact with other people. Since the late 1800s, the argument regarding how the presence of person/people affects motor or cognitive performance compared with a solo condition has continued [16, 17]. In recent years, detailed investigation of the effect of social facilitation [17] has been conducted using more detailed experimental settings. For example, it has been shown that the observing movements of others induces synchronization in one’s movement speed during a competitive game [18], facilitates movement adaptation [19], and influences the prediction of another individual’s movement [20]. Although risk-seeking behavior has been reported in motor tasks in which subjects perform tasks alone, the presence of other people may influence sub-optimal motor decisions.

Here, we investigated how humans alter their motor decision-making in a competitive game (a variant of the “chicken game”), which requires naturalistic interactions with other people. We had two main hypotheses. First, if the decision system simply imitates an opponent’s movement, then a linear relationship between the subject’s and opponent’s decisions should be observed. If this is correct, optimal decisions should be achieved when the opponent’s decisions are also optimal. This hypothesis is based on the evidence that an unintended imitation of movement speed or distance occurs in a competitive situation [18]. Second, if the decision system adaptively adjusts the motor plan according to the opponent’s movements, then a non-linear relationship between the subject’s and opponent’s decisions should be observed. If this hypothesis is true, optimal decisions should be achieved when the opponent’s decisions are sub-optimal.

To test these hypotheses, we assessed subjects’ behavior during competition with a virtual opponent who behaved either optimally or sub-optimally. First, we show that the direction of the sub-optimality of motor decisions is reversed from risk-seeking to risk-averse at the beginning of a competitive situation. Second, following this reversal of sub-optimality, we demonstrate that competition with sub-optimal risk-averse opponents promotes optimal decision-making. Finally, to explain these findings, we confirm that the subjects’ decisions are affected by opponents’ decisions in a non-linear function.

## Results

Subjects performed a quick out-and-back reaching task (moving forward from the start position and returning to the start) using a pen-tablet (Fig 1A and B). A cursor corresponding to the position of a digitized pen was presented on a vertical screen. The endpoint of each movement was defined as the maximum y-position (Fig 1B), and the subjects were rewarded depending on the endpoint following an asymmetric gain function in each trial (Fig 1C). The subjects scored more points if the endpoint was located closer to a green boundary line (set 30 cm forward from the start position); however, if the boundary line was crossed, the score was set at 0. The nature of this game resembles several situations in sports, such as a penalty kick in soccer, service in tennis, the strike zone in baseball, or take-off in ski jumping. The use of this asymmetric gain function is validated by the fact that this function could reveal a sub-optimal risk-seeking or risk-averse behavior [4]. The subjects could aim at any point on the screen. For the selected aim point, the actual endpoint was probabilistic due to the inherent noise of the motor system. Therefore, the subjects were required to make motor decisions regarding where to aim while considering this inherent motor noise.

Figure 2 describes the experimental protocols for five different experiments. Experiment 1 comprised three tasks (training, individual, and competitive tasks) in three experimental sessions. Descriptions of the training task can be found in *Experimental task*. The individual task required the subjects to maximize the total score within each block (10 trials/block). The competitive task required them to perform a trial alternately with their opponents and to achieve a higher total score than their opponents within each block (Fig. 1D). The subjects were randomly divided into three subgroups: *risk-neutral* (Experiment 1a), *risk-averse* (Experiment 1b), and *practice* (Experiment 1c) groups. As shown in Figure 2, the subjects in the risk-neutral and risk-averse groups performed 5 blocks of the individual task (baseline), 12 blocks of the competitive task (competition), and 5 blocks of the individual task (washout). In the practice group, the subjects performed 5 blocks of the individual task (baseline), again 17 blocks of the individual task (individual), and 4 blocks of the competitive task (competition).

In the risk-neutral group, the subjects (N = 9) competed against virtual opponents whose aim points were set at the optimal aim point (see *Experiment 1*). The optimal aim point was calculated by maximizing the expected gain based on each subject’s endpoint variability over the past 40 trials before starting each block of the competitive task (see *Model assumptions*). Because the subjects’ endpoint variability decreased with the progression of the block, risk-neutral opponents’ aim points gradually increased (red line in Fig 3A). In the risk-averse group, the subjects (N = 8) competed against the opponents who gradually changed their aim point from optimal to sub-optimal and risk-averse (red line in Fig 3B). The opponents’ actual endpoint varied from trial to trial and followed a Gaussian distribution. To distinguish the effect of the opponents, the subjects (N = 10) in the practice group continued the individual task (Fig. 3C).

Based on the Bayesian decision theory [21–23], we determined each subject’s risk-sensitivity in the individual (baseline) and competitive tasks as the deviation of the actual aim point (observed mean endpoint) from the optimal aim point (see, *Model assumptions*). If the actual aim point was larger than the optimal aim point (i.e., a positive value), it indicated that the subject adopted a sub-optimal, risk-seeking strategy (seeking a high one-trial reward with a high probability of failure). In contrast, if the actual aim point was smaller than the optimal aim point (i.e., a negative value), it indicated the adoption of a sub-optimal, risk-averse strategy (seeking a low one-trial reward avoiding high probability of failure). If risk-sensitivity was close to 0, the subject was considered to have made optimal, risk-neutral decisions.

Our primary purpose was to investigate what factors in the competition influenced a sub-optimal motor plan taken in the individual task, and how they did so. Before the start of the competition, we did not provide the subjects with any information about their opponents, which might have changed their behavior over the short term. Therefore, we first focused on the time-series of reaching endpoint from the baseline to competition. Later, to describe the effects of opponents over the longer term, we focused on the time-series of risk-sensitivity and aim point (mean endpoint) in each block of the competition. Fig 4A and B illustrate the time series of the reaching endpoint from the baseline to competition. A comparison of the actual and optimal aim points revealed that the subjects adopted a risk-seeking strategy at the baseline in both the risk-neutral group (Fig 4A’; two-tailed paired-sample *t*-test: *t* [8] = 4.65, *p* = 0.002, *d* = 1.29, mean difference = 0.68, 95% CI = [0.35, 1.02]) and the risk-averse group (Fig 4B’; *t* [7] = 5.03, *p* = 0.002, *d* = 2.60, mean difference = 1.15, 95% CI = [0.61, 1.69]). However, they shifted their strategy to be risk-averse by decreasing the reaching endpoint from the baseline at the beginning of the competition (Fig 4A, A’, B, and B’). The average endpoint from the first to fifth trials after the competitive task started significantly decreased from the observed mean endpoint (actual aim point) at the baseline in the risk-neutral group (Fig 4A’; two-tailed paired-sample *t*-test: *t* [8] = 2.76, *p* = 0.025, *d* = 1.12, mean difference = 0.79, 95% CI = [0.13, 1.44]) and the risk-averse group (Fig 4B’; *t* [7] = 3.06, *p* = 0.018, *d* = 1.30, mean difference = 1.06, 95% CI = [0.24, 1.87]). This effect was seen neither when the subjects in the practice group continued the individual task (Fig 4C and C’; *t* [9] = 0.49, *p* = 0.63, *d* = 0.18, mean difference = 0.15, 95% CI = [−0.55, 0.85]), nor when the subjects in the risk-neutral and risk-averse groups moved from competition to washout (Supplementary Fig 1A, A’, B, and B’), nor after the second block of the competition (Supplementary Fig 2). A group level comparison [24] revealed a significant interaction between the experimental group and experimental session (Supplementary Fig 3A). Although the risk-seeking strategy was robust even after repetitive practice for 9 days in a similar experimental setting [6], our results indicated that it could be switched to a risk-averse strategy when competing with a new opponent. In other words, interactions with other people significantly altered motor decision-making.

We further validated this reversal of risk-sensitivity in Experiment 2 and 3. Since human motor control is influenced by both intrinsic uncertainty of the motor system [13] and extrinsic uncertainty of the environment [25–27], we attempted to attenuate these uncertainties before starting the competition. The subjects in the practice group from Experiment 1c performed the competitive task with risk-neutral opponents after completing 22 blocks of the individual task (Fig 2). This practice reduced intrinsic uncertainty; the standard deviation of the reaching endpoint for the last five blocks was 0.73 times lower than that for the first five blocks. In Experiment 2, a new group of 11 subjects was recruited (presentation group). The subjects first completed five blocks of the individual task (baseline). To attenuate the extrinsic uncertainty of the opponents’ behavior, the subjects were then shown the movement of the risk-neutral opponents for 10 trials prior to starting the competitive task (red line in Fig 4E, Fig.2). Although the subjects improved reaching accuracy or acquired knowledge of their opponents in advance, the reversal of risk-sensitivity (from risk-seeking to risk-averse) occurred at the beginning of the competitive task (Fig 4D, D’, E, and E’). In the practice group, the average endpoint from the first to fifth trials of the first competitive block significantly decreased from the observed mean endpoint in the last 50 trials before the competitive task started (Fig 4D’, two-tailed paired-sample *t*-test: *t* [9] = 5.68, *p* = 0.0003, *d* = 2.18, mean difference = 1.07, 95% CI = [0.65, 1.50]). In the presentation group, the average endpoint in 1st–5th trials of the first competitive block significantly decreased from the baseline mean endpoint over 50 trials (Fig 4E’, *t* [10] = 5.29, *p* = 0.0004, *d* = 0.94, mean difference = 0.95, 95% CI = [0.55, 1.35]).

We hypothesized this bias to be triggered by virtual competition with the computer opponent. To test this, we recruited 14 pairs of subjects (human vs. human group) for Experiment 3. A (preceding) subject first performed the first trial of the competitive task, and then, the other (following) subject performed the first trial. Both subjects were instructed to achieve a higher total score in 10 trials (12 blocks in total) than their opponent (see *Experiment 3*). When two subjects competed with each other, a similar trend (decrease of endpoint from the baseline) was observed (Fig 4F and F’ and Supplementary Fig. 4A and A’). The average endpoint in 1st–5th trials of the first competitive block significantly decreased from the baseline mean endpoint in the preceding subject (Supplementary Fig 4A’; two-tailed paired-sample *t*-test: *t* [13] = 2.89, *p* = 0.013, *d* = 1.03, mean difference = 0.67, 95% CI = [0.17, 1.18]) and the following subject (Fig 4F’; *t* [13] = 5.86, *p* = 0.0001, *d* = 2.01, mean difference = 1.17, 95% CI = [0.74, 1.60]). Regarding the effects of practice, observation, and human opponent on the decrease of the endpoint, there were no significant interactions between the experimental group and experimental session (Supplementary Fig 3B, C, and D). Taken together, these findings suggest a clear tendency to abandon an original risk-seeking strategy and start a competition in a conservative manner even when the intrinsic and extrinsic uncertainties are attenuated or when competing against a human opponent.

Following the reversal of risk-sensitivity at the onset of the competitive task, we investigated the influence of the opponents’ decision-making on the subjects’ risk-sensitivity. Again, the risk-seeking strategy (positive value of risk-sensitivity) was adopted at the baseline in the three groups (Fig 5), which remained the same in the practice group. In the practice group, the risk-sensitivity from the 8th to 12th blocks of the individual task was significantly larger than 0 (Fig 5 magenta; two-tailed one-sample *t*-test from 0: *t* [9] = 6.68, *p* = 0.0001, *d* = 3.15, mean difference = 0.88, 95% CI = [0.58, 1.18]). In the risk-neutral group, the strategy was partly modulated (Fig 5 green), but the risk-sensitivity in the last 50 trials of the competitive task (from the 8th to 12th block) remained significant (Fig 5 green; two-tailed one-sample *t*-test from 0: *t* [8] = 4.06, *p* = 0.004, *d* = 2.03, mean difference = 0.56, 95% CI = [0.24, 0.88]). However, when the opponent was risk-averse, the optimal risk-neutral strategy was achieved (Fig 5 blue). The risk-sensitivity in the last 50 trials of the competitive task (from the 8th to 12th block) was not significantly different from 0 (Fig 5 blue; two-tailed one-sample *t*-test from 0: *t* [7] = −0.03, *p* = 0.97, *d* = −0.02, mean difference = −0.01, 95% CI = [−0.59, 0.57]).

Furthermore, two-way mixed design ANOVA revealed a significant group [3] × block [12] interaction (Supplementary Fig 5A; *F* [22, 264] = 2.12, *p* = 0.002, *η*^2^ = 0.09) and a significant group [3] × session [2] interaction (Supplementary Fig 5B; *F* [2, 24] = 10.44, *p* = 0.001, *η*^2^ = 0.12). No significant differences were observed among the three groups in terms of the standard deviation of the reaching endpoint or optimal aim point (Supplementary Fig 6A and B). These results indicate that the sub-optimal risk-seeking strategy was modified by the presence of the opponent. Specifically, the optimal risk-neutral strategy was promoted by competition with a sub-optimal risk-averse opponent.

Next, we addressed why the competition with sub-optimal risk-averse opponents led to optimal and risk-neutral decision-making. To specify the relationship between the opponents’ and subjects’ decisions, we calculated the following indices as the measures of motor decision-making: *A*_*i*_, mean endpoint across five blocks in the individual task, *A*_*c*_, mean endpoint across each block in the competitive task and *A*_*o*_, opponents’ mean endpoint across each block in the competitive task. The subjects in the risk-neutral group gradually increased their aim point as the opponents’ aim point increased (Fig 3A; correlation *r* between red and blue bold lines = 0.82, *p* = 0.001). In contrast, there was no such correlation in the risk-averse group, and the subjects maintained their aim point even though the opponents’ aim point decreased (Fig 3B; correlation *r* between red and blue bold lines = 0.45, *p* = 0.15). When repeating the individual task, a significant increase in the aim point was observed over 12 blocks (Fig 3C; main effect of the block in one-way within-subject ANOVA: *F* [11, 99] = 2.74, *p* = 0.004, *η*^2^ = 0.23). To further explore this relationship, Fig 6A plotted the subjects’ relative aim points (defined as *A*_*c*_ − *A*_*i*_) in the two competitive groups against the opponents’ relative aim points (defined as *A*_*o*_ − *A*_*i*_). If the opponents’ decisions linearly influenced the subjects’ decisions, the slopes of regression line across the right and left halves of the plot should be similar. If this linear relationship is valid, the subjects should make optimal decisions when the opponents make optimal decisions. In contrast, if the opponents’ decisions non-linearly influence the subjects’ decisions, the slopes should be different. If this non-linear relationship is valid, the subjects should make optimal decisions when the opponents make sub-optimal decisions. As shown in Fig 6A, we found a gentler slope in the left half of the plot than in the right half. To assess the statistical difference, we calculated the slopes of regression lines in bootstrapped samples (Fig 6B). The mean slopes were 0.21 (95% CI = [0.07, 0.35]) for the left half of the plot and 0.63 (95% CI = [0.41, 0.83]) for the right half of the plot, that were significantly different (Fig 6B; permutation test: *p* < 0.001). We also fit the overall data in Fig 6A using linear (*Y* = *β*_0_ ∗ *X* + *β*_1_) and quadratic (*Y* = *β*_0_ ∗ *X* + *β*_1_ ∗ *X*^2^ + *β*_2_) models and found that the quadratic model fit the data better than the linear model (Linear model: Akaike information criteria [AIC] = 1151.7, Bayesian information criteria [BIC] = 1158.2, AIC for small sample size [AICc] = 1151.8, Adjusted R^2^ = 40.7%; Quadratic model: AIC = 1144.3, BIC = 1154.0, AICc = 1144.4, Adjusted R^2^ = 43.3%). Overall, these results suggest that the subjects’ decision-making was influenced by a non-linear function of the opponents’ decision-making—competition with sub-optimal, risk-averse opponents led to optimal decision-making—.

One concern regarding our study is that the individual task and competitive task had different goals, which makes a difference in the optimal strategy between tasks. However, maximizing one’s own total score could also be a near-optimal strategy in the competitive task because it requires one to compete for a magnitude of the total score. To visualize this, we calculated the chance of winning (color bar in Fig. 6A, see *Simulation of the chance of winning*). When the opponent is risk-neutral (the value on the X-axis is around −1 to 1), the region of a higher chance of winning is limited. In contrast, for the risk-averse opponent (the value on the X-axis is below −1), this region extends triangularly. This means that the strategy to win against the risk-averse opponent is redundant. The actual data in the risk-averse group (white circle) were distributed within the optimal region which maximizes the chance of winning. Therefore, the strategy taken by the risk-averse group can be interpreted as the optimal strategy in terms of maximizing not only their own total scores but also their chance of winning. Notably, the data were fit better with a non-linear function of opponents’ aim points than a linear function. If the data (white circle) followed a linear function of opponents’ aim points, this linear strategy could maximize the chance of winning but would fail to maximize the expected reward because the strategy turned to be risk-averse. Thus, the non-linear strategy can be the only way to achieve the optimality in terms of maximizing the total scores and chance of winning.

We validated the achievement of optimal motor decision-making in Experiment 4. The non-linear relationship between the subject’s and opponent’s decisions predicts that the subjects maintain their aim point even when the opponent’s aim point is considerably shorter. Therefore, if the non-linear relationship is valid, optimal decisions should be achieved also when the opponent’s decisions are highly sub-optimal. To examine this hypothesis, a new group of six subjects was recruited for Experiment 4 (highly risk-averse group). The subjects competed with a highly risk-averse opponent who aimed further from the boundary line from the initial to last block of the competitive task (see red line in Fig 7B). Similar to the risk-averse group in Experiment 1b, the subject’s risk-sensitivity was modulated (Fig 7A). Although the subjects showed a risk-seeking strategy at the baseline, there was no significant difference between the subject’s risk-sensitivity and the risk-neutral value (i.e., 0) in the last 50 trials (blocks 8–12) of the competitive task (Fig 7A; two-tailed one-sample *t*-test from 0: *t* [5] = −0.26, *p* = 0.80, *d* = −0.11, mean difference = −0.01, 95% CI = [−1.1, 0.86]), suggesting that the optimal risk-neutral strategy was achieved. We replicated a non-linear relationship between the opponents’ and subjects’ decisions. The subjects decreased their aim point from the baseline and maintained it, whereas the highly risk-averse opponents consistently aimed further from the boundary line (Fig 7B). The subjects’ relative aim points from the baseline were non-linearly influenced as a function of the opponents’ relative aim points (Fig 7C). These results demonstrate the validity of a non-linear relationship between the subjects’ and opponents’ decisions and that a sub-optimal risk-averse opponent promotes optimal and risk-neutral motor decision-making.

We demonstrated a clear effect of the competition on motor decision-making. However, there is a possibility that components underlying the competition led to this effect. The competitive task used in this study had two components: observing the opponent’s performance and exceeding the opponent’s total score. Could the behavioral changes be solely derived by observing the opponent’s performance or attempting to exceed the total score in a situation not involving the opponent? To investigate this question, new groups of six subjects were recruited for Experiment 5a (observation group) and Experiment 5b (threshold group). We prepared new tasks by modifying the previous tasks. In the observation task, the subjects were shown the movement trajectory, movement endpoint, score in a trial, and total score of the risk-averse opponent, as in the competitive task. However, they were required to maximize the total score. In the threshold task, there was no opponent as in the individual task. However, the subjects were required to exceed a total score which was presented at the beginning of each experimental block. The threshold of the required total score was set as the score that the risk-averse opponent would achieve. Figure 8A and B illustrate the actual aim point (observed mean endpoint) in the observation group and threshold group, respectively. We did not find an inhibition of the aim point as shown in Fig 3B. In the observation group, the subject’s aim point at the baseline (mean ± SD = 27.92 ± 0.92 cm) was not significantly different from that of the last 50 trials of the observation task (mean ± SD = 27.85 ± 0.70 cm) (two-tailed paired-sample *t*-test: *t* [5] = 0.40, *p* = 0.71, *d* = 0.16, mean difference = 0.07, 95% CI = [−0.40, 0.55]). Similarly, in the threshold group, the subject’s aim point at the baseline (mean ± SD = 27.87 ± 0.45 cm) was not significantly different from that of the last 50 trials of the threshold task (mean ± SD = 27.64 ± 1.00 cm) (two-tailed paired-sample *t*-test: *t* [5] = 0.45, *p* = 0.67, *d* = 0.18, mean difference = 0.23, 95% CI = [−1.08, 1.53]). Therefore, we verified that the competition, rather than solely observing the opponent’s performance or attempting to exceed the total score, promotes optimal motor decision-making. The behavioral changes could be derived by an awareness of the competition [28] induced by the combination of these components.

One confounding factor was how to display the opponent’s score. In the competitive task, the subjects were provided with their opponent’s score in each trial. In contrast, in the threshold task, the subjects were provided with the total score they should attempt to exceed at the beginning of each block. This difference raises the question of whether the subjects would show optimal motor decision-making if they were asked to exceed the expected score in each trial instead of the total score at the end of the block. Clearly, however, the subjects in the threshold task could easily calculate how many points they needed in each trial by simply dividing the required total score by 10 trials. We therefore expected that this difference in the experimental settings would be slight.

## Discussion

For over a decade, sub-optimal and risk-seeking behaviors have been repeatedly confirmed in studies of motor decision-making tasks with an asymmetric gain function [4–6, 10], which require a choice with different variances of pay-off [2, 3, 9, 11] and involve a speed-accuracy trade-off [1]. Despite such findings, solutions to promote optimal motor decision-making are lacking. Here, we assessed the potential effect of interaction with opponents on sub-optimal motor decision-making with a prediction that other people’s actions/intentions can influence the subjects’ motor system [18, 18, 29, 30]. First, we found that the subjects’ risk-seeking strategy in the individual task reversed to risk-averse strategy at the very beginning of the competitive task (Fig 4). Second, optimal motor decisions were promoted by competition with a risk-averse opponent (Fig 5). This optimal decision-making was induced by a non-linear influence of the opponents’ decisions (Fig 6).

The reversal of risk-sensitivity was robustly shown through several experiments (Fig 4). However, this switching of strategy from the individual task is not convincing. In the individual task, the subjects were instructed to maximize the total score. At the beginning of the competitive task, when the subjects did not know the opponent’s strategy, they should have maintained their original strategy to maximize the total score and beat the opponent. The data showed large decrease in the endpoint in the first trial, which recovered thereafter (Fig 4). This might reflect an exploration of the new task, but a simple random exploration does not explain the decrease in the endpoint. In other words, if it reflected only an exploration, the endpoints would have either increased or decreased. A possible explanation for this behavior is that the subjects sought a better strategy believing that they would compete against a weak opponent who aimed for a lower score. If the subjects believed that the opponent was strong and would aim for a higher score, they would not have changed their original strategy. Therefore, this amount of decrease reflects the subjects’ risk-premium [31] that they would recover the points in later trials even if they scored fewer points at the beginning. By sacrificing the cost of scoring fewer points, the subjects may be seeking an optimal strategy to beat a weak opponent.

We also found that the subjects’ motor decisions were non-linearly influenced by the opponents’ decisions (Figs 3&6). The subjects increased their aim point when the opponents also aimed for a higher score (Fig 3A). In contrast, when the opponents aimed for a lower score, the subjects did not change their aim point (Fig 3B). Therefore, the subjects adaptively altered their decisions according to the opponents’ decision, rather than imitating it. If the opponents’ decision linearly affected the subjects’ decision and imitation occurred, the subjects would have also aimed for a lower score when the risk-averse opponent decreased the aim point. The decision strategy that the subjects adopted can be interpreted as a variant of the win-stay lose-shift strategy [32]. Importantly, in terms of the win-stay part (Fig 3B), the subjects decreased their aim point from the individual task and then let the strategy “stay”, rather than adopting the original risk-seeking strategy and then letting the risk-seeking strategy “stay”. If the subjects simply would have simply adopted the win-stay lose-shift strategy, they would kept letting the original risk-seeking strategy stay when competing with the highly risk-averse opponents (Experiment 4) because they rarely lost to these very weak opponents. Therefore, these results suggest that the opponents had both the inhibitory effect and non-linear effect on the subjects’ motor decision-making and that the mixture of these effects induced the optimal and risk-neutral strategy. The modulation of the competition-induced risk-sensitivity was retained even in a situation not involving the opponents after the competition in the highly risk-averse group (Supplementary Fig 7). Further research is necessary to clarify how the opponent was modeled into the decision system to generate an optimal motor plan.

Strategic decision-making has been investigated in game theory tasks that require players to make discrete choices [33]. In the Prisoner’s Dilemma game [34]—a standard game theory task—two prisoners have two choices, cooperation or defection, which determine four possible pay-offs (prison sentences). In the current study, however, the subjects decided where to aim to beat their opponent. Such continuous choice (motor decision-making) is often required in competitive sports (soccer, tennis, baseball, golf, darts, ski jumping etc.). Therefore, the current study highlighted the characteristics of movement strategy in competitive situations. Specifically, we clarified how interaction with opponents improved sub-optimal motor decision-making. When humans practice a motor task alone (without opponents), repetitive practice has been shown to improve movement accuracy but not movement strategy [6]. Our findings suggest that competition with an opponent, particularly a risk-averse opponent, is an effective means to promote an optimal and risk-neutral movement strategy. In behavioral economics, Richard Thaler (2017 Nobel economics winner) proposed the term “nudge” to be a means of behavioral change as human decision-making is systematically biased under bounded rationality (nudge refers to the choice architecture which guides an individual’s choice towards a beneficial one while maintaining freedom of choice) [35]. Presence of people can be interpreted as one of the nudges which can alter sub-optimal motor choice. This information may be helpful for sports trainers and coaches to achieve a better motor performance—the importance of other people should be considered in developing a training protocol in sports.

## Methods

### Subjects

We recruited 84 healthy adults (52 males; 20.4 ± 2.0 years) for the experiments. All experiments were conducted following a random assignment of the study subjects. This study was approved by the ethics committee of Tokyo University of Agriculture and Technology and was carried out in accordance with the approved guidance. The subjects provided written informed consent and were unaware of the purpose of the experiment.

### Apparatus

We used a pen-tablet with sufficient workspace to measure the subjects’ arm-reach movement (Wacom, Saitama, Japan, Intuos 4 Extra Large; workspace: 488 × 305 mm). The subjects made a quick out-and-back reaching movement holding the digitized pen on the pen-tablet (Fig 1A). The position of the digitized pen was sampled at ~144 Hz with a spatial resolution of 0.01 mm. The subjects manipulated a cursor on a vertical screen whose position was transformed from the pen position with a maximum delay of 6.9 ms (Asus, Taipei, Taiwan, VG-248QE; size: 24 inches, refresh rate: ~144 Hz). The scale of the pen and cursor position was 1:1. All stimuli were controlled using Psychophysics Toolbox [36, 37].

### Experimental task

There were five tasks: *training*, *individual*, *competitive*, *observation*, and *threshold tasks*. For all tasks, to begin each trial, subjects moved a blue cursor (radius: 0.3 cm) to a white starting position (radius: 0.4 cm) presented on the vertical screen. After a 1-s delay, a horizontal white line (width: 0.1 cm) appeared 30 cm from the starting position and turned green after random intervals of 0.8–1.2 s, indicating a “go” signal. In this paper, this green line is referred to as the “boundary line”. After the “go” signal, the subjects made a quick out-and-back reaching movement to rapidly move the cursor forward and then return it below the starting position. The subjects received an online feedback about how the cursor was moving but could not see their own hand position since it was covered with a box. We recorded the endpoint of each movement as the maximum y-position (Fig 1B). If the subjects did not return the cursor within 600 ms (time-out), a message stating “Time-out. More quickly!” was presented with a warning tone. If the subjects successfully completed the trial, a yellow cursor (radius: 0.3 cm) appeared at the position of the reaching endpoint for 2 s. After the feedback period, the subjects proceeded to the next trial.

#### Training task

Before the individual task, a training task was assigned to allow the subjects to practice the reaching movement. The subjects were required to reach the green boundary line. After each movement, if the yellow cursor overlapped with the green boundary line, the message “Hit!” appeared on the screen with a pleasant sound. The training task comprised 50 trials.

#### Individual task

In the individual task, the subjects were awarded points depending on the reaching endpoint (Fig 1C). More points were awarded when the endpoint was closer to the green boundary line at 30 cm; however, the score for this trial fell to 0 if the endpoint crossed the boundary line. When a mistrial occurred, a “Miss!” message appeared on the screen with a flashing red lamp along with an unpleasant alarm. Of note, 0 points were also awarded if the endpoint was within 7 cm from the start position, but no such trials were observed. In case of time-out, 0 points were awarded. In the feedback period, the current and total points were presented along with the reaching endpoint. The subjects were instructed to maximize the total points in each experimental block which comprised 10 trials each.

The use of an asymmetric gain function is validated by the fact that this function could reveal a sub-optimal risk-seeking or risk-averse behavior [4]. If a function with gain distributed symmetrically around the boundary line was used, we could not have measured the subjects’ risk-sensitivity because aiming the boundary line was obviously optimal for all subjects.

#### Competitive task

The competitive task was performed against a computer or human opponent (Fig. 1D). Each experimental block comprised 10 trials. The subjects performed the reaching movement in the same way as in the individual task. After the feedback period, a red cursor (radius: 0.3 cm) was shown on the screen, indicating the opponent’s turn. In the competitive task with the computer opponent, the opponent’s cursor movement (trajectory) was automatically manipulated based on pre-recorded sample trajectories made by the experimenter. Each movement endpoint was determined by the pre-programmed algorithm described below (Manipulation of computer opponent). In the competitive task, the subjects were instructed to win the game by achieving a higher total score than their opponents at the end of each experimental block.

#### Observation task

In the observation task, the subjects were required to maximize the total score in the same setting as the competitive task but with modified instructions. In each trial, they were shown the movement trajectory, movement endpoint, score in a trial, and total score of the opponent. However, the task’s goal was not to beat the opponent but to obtain the highest total score. At the end of each experimental block comprising 10 trials each, their total score was displayed.

#### Threshold task

In the threshold task, the subjects were required to exceed the presented total score in the same setting as the individual task except for different instructions. This task did not involve an opponent, and its goal was not to maximize the total score but to exceed a required total score which was presented at the beginning of each experimental block. Each block comprised 10 trials. At the end of each block, the subjects received a binary feedback depending on whether they exceeded the required total score or not, indicating they won or lost, respectively.

#### Manipulation of computer opponent

For the trials involving a computer opponent, we randomly sampled the endpoint of each trial from a Gaussian distribution with mean *αE*^∗^ and variance *σ*^2^, where *α* represents the coefficient that determines the opponent’s risk-sensitivity and *E*^∗^ represents the optimal mean endpoint maximizing the expected reward given the variance of reaching endpoint *σ*^2^. Before each experimental block in the competitive task, we determined the value of *σ*^2^ by calculating the subject’s reaching variance over the past 40 trials. This means that the computer opponent always had the same reaching accuracy as the subject. We then defined the computer’s mean endpoint as *αE*^∗^. We could dynamically manipulate the endpoint of the opponent by changing the coefficient *α*.

### Experiment 1

Experiment 1 comprised three subgroups with 9 (3 males; 20.4 ± 2.7 years), 8 (5 males; 19.1 ± 0.6 years), and 10 (7 males; 19.9 ± 2.0 years) subjects assigned to the *risk-neutral* (Experiment 1a), *risk-averse* (Experiment 1b), and *practice* (Experiment 1c) groups, respectively. For the risk-neutral and risk-averse groups, there were three experimental sessions: *baseline* comprising 5 blocks of the individual task, *competition* comprising 12 blocks of the competitive task, and *washout* comprising 5 blocks of the individual task. For the practice group, there were baseline (5 blocks of the individual task) and *individual* (17 blocks of the individual task) sessions. Following 22 blocks of the individual task, the subjects completed 4 blocks of the competitive task. See Figure 2 for experimental protocols.

As manipulation of the computer opponent, the value of *α* for the risk-neutral opponent was set as *α* = 1 for all 12 experimental blocks, i.e., the computer opponent always behaved as an optimal risk-neutral decision-maker who gradually aimed closer to the boundary line along with the reduction of the reaching variance. For the risk-averse opponent, this was set as *α* = 1 for blocks 1–4, decreased in steps of 0.15 for blocks 5–8, and finally set as *α* = 0.925 for blocks 9–12, i.e., the computer opponent behaved as a sub-optimal risk-averse decision-maker who gradually aimed further from the boundary line. The differences in movements between the two computer opponents can be seen in Fig 3A and B. Note that both opponents were set as *α* = 1 for the first four blocks. For four blocks of the competitive task in the practice group, the value of a was set as *α* = 1.

### Experiment 2

Experiment 2 was conducted to investigate whether the extrinsic uncertainty of the opponents’ behavior affected the subjects’ risk-averse strategy at the beginning of the competitive task. Eleven subjects (10 males; 20.2 ± 2.1 years) were recruited (*presentation* group). The experimental protocol was the same as Experiment 1a and b except for a *presentation* session between the baseline and the competition (Fig 2). Prior to the first block of the competitive task, the subjects observed the movement of the risk-neutral opponent (*α* = 1) for 10 trials. In the competition, the value of a was set as *α* = 1 for all 12 experimental blocks.

### Experiment 3

We conducted Experiment 3 to see whether the risk-averse strategy at the beginning of the competitive task was triggered by the virtual competition with the computer opponent. Fourteen pairs of subjects (16 males, 12 females; 21.2 ± 2.0 years) were recruited (*human vs. human* group). Two sets of pen-tablet and screen were prepared. A vertical partition was used to separate the two subjects, preventing verbal and non-verbal communication during the experiment. There were baseline comprising 5 blocks of the individual task and competition comprising 12 blocks of the competitive task (Fig 2). Two subjects alternatively performed each block of the individual task. The screen of each subject was turned off while the other subject was performing the individual task to prevent them from seeing each other’s performances. In the competitive task, the same stimuli were presented on each screen. Each subject performed the task alternatively from trial to trial. They were instructed to achieve a higher total score than the other subjects at the end of each experimental block.

Because we focused on the reversal of risk-sensitivity from the individual to competitive tasks, we analyzed the very first block of the competitive task (Fig 4). In all risk-neutral, risk-averse, practice, and presentation groups, the value of a at the first block was controlled to be *α* = 1, which facilitated examination of the effect of practicing the task or observing the opponent performance on the reversal of risk-sensitivity. For a similar reason, we analyzed the first block of the competitive task in the human vs. human group.

### Experiment 4

We conducted a follow-up experiment to validate the achievement of risk-neutrality by the competition with a risk-averse opponent. To do so, we induced a computer opponent to display a highly risk-averse behavior. Six subjects (4 males; 19.7 ± 1.5 years) were recruited in the *highly risk-averse* group. The same experimental protocol as Experiment 1a and b was used (Fig 2). In Experiment 4, we set the coefficient as *α* = 0.925 for all (1–12) blocks of the competitive task to ensure the opponent exhibited a highly risk-averse behavior from the initial block of the competition. Further, the variance of the opponent’s reaching endpoint *σ*^2^ was always set as the subject’s reaching variance in the last 40 trials of the baseline (blocks 2–5), because we attempted to have the computer opponent consistently aim further from the boundary line. The movement of the opponent can be seen in Fig 7B.

The sample size was determined by *a priori* power analysis using G*power [38]. For the determination of the effect size of two paired samples [38], we used the subjects’ data in Experiment 1b. Because the competition with the risk-averse opponent induced the optimal and risk-averse strategy, we compared the subject’s mean risk-sensitivity at the baseline (mean ± standard deviation = 1.15 ± 0.65 cm) with that in the last 50 trials (blocks 8–12) of the competition (mean ± SD = −0.01 ± 0.70 cm). As a result, we obtained the effect size of *d* = 1.54 (correlation between two samples *r* = 0.38). The *a priori* power analysis (matched pairs *t*-test, *α* = 0.05, *β* = 0.80, *d* = 1.54) provided a sample size of n = 6. We thus recruited six subjects in Experiment 4.

### Experiment 5

Experiment 5 was conducted to see whether the behavioral changes induced by the competition with risk-averse opponent solely resulted from the observation of the opponent’s behavior or the achievement of the opponent’s total score. Two sets of six subjects were recruited in the *observation* group (2 males; 21.0 ± 2.1 years, Experiment 5a) and *threshold* group (5 males; 19.7 ± 0.8 years, Experiment 5b). We determined the sample size using *a priori* power analysis. If observing the opponent’s behavior or exceeding the opponent’s total score influenced the subject’s motor decision-making, we would find the same effect as the risk-averse group in Experiment 1b. We thus determined the sample size to be six in Experiment 5 based on *a priori* power analysis.

For the observation group, there were three experimental sessions: *baseline* comprising 5 blocks of the individual task, *observation* comprising 12 blocks of the observation task, and *washout* comprising 5 blocks of the individual task. For the threshold group, we performed a *threshold* session comprising 12 blocks of the threshold task instead (Fig 2). As a manipulation of the computer opponent in the observation task, we chose the opponent’s risk-sensitivity value a to be the same as that for the risk-averse group (Experiment 1b) to control the experimental setting regarding the opponent. In the threshold task, we set the threshold of total score as the score that the risk-averse opponent would achieve. Before each block begun, we calculated the expected total score that the risk-averse opponent would achieve for given a value. We set this total score as the threshold. The threshold of total score was the highest in blocks 1–4, and it gradually decreased.

### Model assumptions

Based on Bayesian decision theory [21–23], we modeled the optimal mean endpoint by maximizing the expected gain for a given sensory motor variability to quantify the subjects’ risk-sensitivity and define the computer opponents’ endpoint. In this model, the expected gain *EG(E)* for a selected aim point (mean endpoint) *E* can be calculated by integrating the gain function *G(e)* with the probability distribution of the movement endpoint *P(e|E)*.

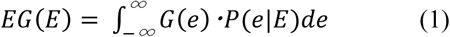

We assumed that the actual movement endpoint *e* is distributed around a selected aim point *E* with sensory motor variability *σ*^2^ according to Gaussian distribution.

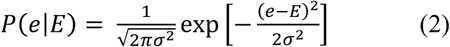

Given a subject’s variability in movement endpoint *σ*^2^, we estimated the optimal mean endpoint by maximizing the expected gain function.

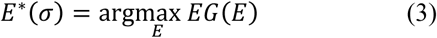

For a detailed description of the model assumption, see Ota et al [5, 6].

We simplified our model based on two assumptions: (1) a subject’s aim point was fixed between trials and (2) the reaching variability remained constant, regardless of the magnitude of the aim point. The first assumption was validated because the reaching variability did not differ depending on whether the aim point was fixed or not (Supplementary Fig 8). The second assumption was validated because the reaching variability did not differ among the risk-neutral, risk-averse, and practice groups (Supplementary Fig 6A), although the higher value of aim point was adopted in the practice group (Fig 3C) and the slower movement duration was taken in the risk-neutral group (Supplementary Fig 9). Further, in our previous study, we confirmed a marginal difference between the model prediction assuming the constant variance and proportional variance to the aim point [6].

### Statistical analysis

To see whether the actual aim point (observed mean endpoint) differed from the optimal aim point at the baseline, we performed a two-tailed paired-sample *t*-test (Fig 4). In Fig 3A’,B’,C’,E’, and F’, the actual and optimal aim point were calculated based on the baseline data (50 trials). In Fig 4D’, these variables were calculated based on the data of the last 50 trials before the beginning of the competitive task. To investigate whether the endpoint decreased at the beginning of the competitive task, we performed a two-tailed paired-sample *t*-test between the average endpoint from the first to fifth trials at the first block of the competitive task and the actual aim point for the baseline data (Fig 4A’, B’, E’, and F’) or the last 50 trials before the competitive task started (Fig 4D’). In Fig. 4, we show that the endpoint was at its lowest at the first trial after the individual task started, and that it gradually recovered up until the sixth trial. We thus pooled the data in the first 5 trials for the statistical analysis. To confirm whether this effect originated from the competition, we compared the average endpoint in 1–5 trials after the individual task restarted with the baseline aim point (Fig 4C’).

In Figure 5, two analyses were performed. At the baseline, we calculated the risk-sensitivity as the difference between the actual and optimal aim points in the baseline data (5 blocks). In the shaded gray area, we calculated the risk-sensitivity as the difference between the actual and optimal aim points in the 8th–12th blocks of the competitive task (risk-neutral and risk-averse groups) or individual task (practice group). We then performed two-tailed one-sample *t*-tests from these risk-sensitivity values to the risk-neutral value (i.e., 0).

To determine whether a slope of regression line differed between the right and left halves of the plot in Figure 6A, we conducted bootstrap sampling and permutation test. For bootstrap sampling, the data of the risk-neutral and risk-averse groups were intermingled and resampled 50,000 times. For each resample, we calculated the slopes of regression line for the right and left halves of the plot (see histograms in Fig 6B). For permutation test, we counted how many times the slope in the left half of the plot was larger than that in the right half to determine *p*-value (null hypothesis was that the frequency would be the same if there is no difference between the slopes of the two halves).

### Simulation of the chance of winning

We performed a Monte Carlo simulation to estimate the chance of winning for an aim point against the opponent’s aim point. Given the opponent’s aim point, variance of the opponent’s endpoint, and variance of the subject’s endpoint in each experimental block, we simulated two samples of 10 trials. One sample was generated from a Gaussian distribution with a simulational aim point (mean) and the variance of subject’s endpoint. The other was generated from a Gaussian distribution with the opponent’s aim point (mean) and variance of the opponent’s endpoint. We simulated these two samples 10,000 times for each aim point. By counting how many times one with a particular aim point could win against an opponent, we estimated the chance of winning as a function of the aim point. Fig 6A shows the average chance of winning over the data in each bin (in steps of 0.25 cm).

## Acknowledgments

We thank Dr. Daichi Nozaki, Dr. Kazutoshi Kudo, Dr. Takuji Hayashi, and Dr. Laurence T Maloney for their inspirational discussions. This research was supported by Grant-Aid for JSPS Fellows No. 17J07822 and Hayao Nakayama Foundation for Science & Technology and Culture No. H29-B-58 awarded to KO and by JSPS KAKENHI No. 18K17894 awarded to KT.

## Author contributions

KO and KT conceived and designed the experiments. KO, MT, KI, and KT performed the experiments. KO and KT analyzed the data and interpreted the results. KO wrote the manuscript and KT revised the manuscript. KO, MT, KI, and KT approved the final manuscript.

## Code availability

The code used to generate the data will be uploaded on a data-sharing website after the manuscript has been officially accepted.

## Data availability

The data will be uploaded on a data-sharing website after the manuscript has been officially accepted.

---

**Fig 1. Experimental set up (“game of chicken”).**

(A) Experimental apparatus. The subjects held a digitized pen on a pen-tablet. The stimuli were shown on a vertical screen in front of the subjects. (B) Trajectory of reaching movement. The subjects made a quick out-and-back reaching movement, by moving forward from the start initial position (white circle) and returning to it. The reaching endpoint (yellow circle) was calculated as the maximum y-position. (C) Asymmetric gain function. The reaching endpoint determined the one-trial score. The maximum score (100 points) was associated with reaching the green boundary line (30 cm). (D) Trial-sequence of the competitive task. The subjects and opponents performed the reaching movement in an alternating order. The current total scores for the subjects and opponents were constantly displayed on the screen, indicating the difference in scores. The reaching trajectories (red and blue) were shown only for clarity.

**Fig 2. Experimental protocol for each experiment.**

The baseline, individual, and washout sessions comprised the individual task, whereas the competition session comprised the competitive task. The observation and threshold sessions comprised the observation and threshold tasks, respectively. Numbers in parentheses denote the number of experimental blocks conducted.

**Fig 3. Effect of opponent’s aim points on subject’s aim points.**

(A–C) Change of aim point (mean endpoint) when competing against a (A) risk-neutral opponent, (B) risk-averse opponent, and (C) performing the individual task. Data is averaged across the subjects, and the shaded area denotes the standard error of the mean. In the risk-neutral group, the subjects’ aim points increased as the opponents’ aim points increased. In contrast, the subjects’ aim points in the risk-averse group did not change, whereas the opponent’s aim points decreased.

**Fig 4. Reversal of strategy from risk-seeking to risk-averse.**

(A–F) Time series of the reaching endpoint in the individual task and the first block of the competitive task. Data is averaged across the subjects and the shaded area denotes the standard error of the mean. The horizontal solid line indicates the observed mean endpoint in the 50 trials of the individual task before the competitive task started (A, B, and D–F) or when the individual task restarted (C). (A’–F’) The bar graphs show the reaching endpoint. Data is averaged across the subjects, and the error bar denotes the standard error of the mean. Opt. indicates the optimal mean endpoint in the individual task for 50 trials, whereas Obs. indicates the observed mean endpoint in the individual task for 50 trials corresponding to the horizontal solid line in (A–F). Further, 1st–5th indicates the average endpoints across the first to fifth trials after the competitive task started (A’, B’, and D’–F’) or when the individual task restarted (C’). * represents p < 0.05, and ** represents p < 0.01 (paired t-test). Open circles denote the data of each subject. In the preceding individual task, the risk-seeking strategy was adopted, indicated by the deviation from the optimal to the observed mean endpoint. However, the decrease in the endpoint was seen at the beginning of the competition (A’ and B’), suggesting that the subjects switched their risk-seeking strategy to risk-averse strategy. This effect was robust even when the competitive task began after the individual task was repeatedly performed (D’), when the opponents’ endpoint was presented in advance (E’), and when the subjects competed against human opponents (F’). When the subjects repeated the individual task, this strategy shift was not observed (C’). The data in the practice group are presented twice (C, C’, D, and D’) because we wanted to know whether the reversal of risk-sensitivity was not derived by performing the individual task (C and C’) and was still observed after a long-term practice of the individual task (D and D’).

**Fig 5. Achievement of risk-neutrality influenced by the opponents.**

Risk-sensitivity values, defined as the difference between the mean endpoint and optimal endpoint, are plotted. BL denotes the risk-sensitivity for 5 blocks of the individual task (baseline). Each block denotes the risk-sensitivity for each block of the competitive task (risk-neutral and risk-averse groups) or for each block of the individual task after the baseline (practice group). The shaded gray area highlights the last five blocks. Data is averaged across subjects, and the shaded area denotes the standard error of the mean. Positive values indicate a risk-seeking strategy, whereas negative values indicate a risk-averse strategy. Green, blue, and magenta asterisks denote *p* < 0.01 from the risk-neutral value (i.e. 0) for the risk-neutral, risk-averse, and practice group, respectively. At the baseline, the risk-seeking strategy was taken in all groups. This trend was consistent in the practice group and partially modulated in the risk-neutral group. In the risk-averse group, the sub-optimal risk-seeking strategy was improved to risk-neutral. Notably, in the risk-averse group, there was no significant difference in the last 50 trials of the competitive task (shaded gray area). Between-group comparison of risk-sensitivity values is shown in Supplementary Fig 5.

**Fig 6. Non-linear influence of opponent’s decisions on subject’s decisions.**

(A) Aim point modulation as a function of the opponents’ relative aim points. The subjects’ relative aim point (mean endpoint in the competitive task *A*_*c*_ − mean endpoint in the individual task *A*_*i*_) is plotted against the opponents’ relative aim point (mean endpoint of the opponent *A*_*o*_ − *A*_*i*_). Black circles indicate the actual data of the risk-neutral group (11[2nd–12th] blocks × 9 subjects), and white circles indicate that of the risk-averse group (11[2nd–12th] blocks × 8 subjects). The slope of regression line in the left (blue) and right (red) halves of the plot predicted the non-linear influence of the opponents. The green dashed line shows a quadratic curve fitted to the data (Y = 0.625X + 0.009X^2^ + 3.17). A color bar denotes the chance of winning when the aim point data falls within a particular region. (B) Histogram of bootstrapped slopes of regression line (50,000 repetition). Vertical lines indicate the mean values of each distribution. Permutation tests revealed a significant difference between the slopes of regression lines, suggesting a smaller influence of the opponents’ decisions on the subjects’ decisions in the left half of the plot than in the right half.

**Fig 7. Competition with a highly risk-averse opponent in Experiment 4.**

(A) Risk-sensitivity value is plotted for the baseline and for each block of the competitive task. A positive value indicates a risk-seeking strategy. * denotes p < 0.05 from the risk-neutral value (i.e., 0). The sub-optimal risk-seeking strategy at the baseline improved during the competition. The shaded gray area denotes the last 50 trials of the competitive task. (B) Change of aim point (mean endpoint) when competing against the highly risk-averse opponent. The opponent aim point was set further away from the boundary line. We replicated the results in the risk-averse group (Fig 3B). The subjects’ aim point decreased from the baseline and retained during the competition. Data is averaged across the subjects, and the shaded area denotes the standard error of the mean. (C) Aim point modulation as a non-linear function of the opponents’ relative aim point. The subjects’ relative aim point (mean endpoint in the competitive task *A*_*c*_ − mean endpoint in the individual task *A*_*i*_) is plotted against the opponents’ relative aim point (mean endpoint of the opponent *A*_*o*_ − *A*_*i*_). Sixty-six data points (11[2nd–12th] blocks × 6 subjects) are plotted.

**Fig 8. Results of the observation and threshold tasks in Experiment 5.**

Change of aim point (mean endpoint) when observing the performance of the risk-averse opponent (A) and attempting to exceed the required total score (B). Data is averaged across the subjects, and the shaded area denotes the standard error of the mean. The horizontal dashed line indicates the observed mean endpoint at the baseline (50 trials). Unlike the competition with the risk-averse opponent, the change of the aim point was not induced solely by observing the opponent’s performance or achieving the required total score.

